# Distinct Sources of a Bovine Blastocyst Digital Image do not Produce the Same Classification by a Previously Trained Software using Artificial Neural Network

**DOI:** 10.1101/424028

**Authors:** Vitória Bertogna Guilherme, Micheli Pronunciate, Priscila Helena dos Santos, Diego de Souza Ciniciato, Maria Beatriz Takahashi, José Celso Rocha, Marcelo Fábio Gouveia Nogueira

**Author notes:** Those authors contributed equally to the study (Vitória Bertogna Guilherme, Micheli Pronunciate, Priscila Helena dos Santos).

## Abstract

We develop an online graphical and intuitive interface connected to a server aiming to facilitate access to professionals worldwide that face problems with bovine blastocysts classification. The interface Blasto3Q (3Q is referred to the three qualities of the blastocyst grading) contains a description of 24 variables that are extracted from the image of the blastocyst and analyzed by three Artificial Neural Networks (ANNs) that classifies the same loaded image. The same embryo (*i.e.*, the biological specimen) was submitted to digital image capture by the control group (inverted microscope with 40x of magnification) and to experimental group (stereomicroscope with maximum of magnification plus 4x zoom from the cell phone). The 36 images obtained from control and experimental groups were uploaded on the Blasto3Q. Each image from both sources was evaluated for segmentation and submitted (only if it could be properly or partially segmented) to the quality grade classification by the three ANNs of the Blasto3Q program. In the group control, all the images were properly segmented, whereas 38.9% (07/18) and 61.1% (11/18) of the images from the experimental group, respectively could not be segmented or were partially segmented. The percentage of agreement was calculated when the same blastocyst was evaluated by the same ANN from the two sources (control and experimental groups). On the 54 potential evaluations of the three ANNs (*i.e.*, 18 images been evaluated by the three networks) from the experimental group only 22.2% agreed with evaluations of the control (12/54). Of the remaining 42 disagreed evaluations from experimental group, 21 were unable to be performed and 21 were wrongly processed when compared with control evaluation.

## 1. Introduction

Brazil is the largest beef exporter in the world with a cattle herd of 218.2 million of animals [1], number that is superior of the Brazilian population (207 million at 2017) [2]. This production is the major economic activity of country, moved US$ 5.3 billion at 2016 and employed 1.6 million people. It is also the major producer of *in vitro* bovine embryos, with 366.517 embryos/year, that is around 70% of the world production [3]. The Brazilian program of *in vitro* production (IVP) of bovine embryos started on 1990, but the first three births only occurred in 1994. This achievement used Nellore breed immature oocytes, frozen-thawed semen and culture system. Currently, the IVP is used commercially for several laboratories to research and multiplication of genetic material in the animal production [4]. This production has utmost importance for international and national improvement in cattle genetics and productivity.

The animal production has undergone many changes in the last 50 years, due to a livestock sector and agricultural research. In this case, the development of *in vitro* techniques for production of cattle embryos has proved to be useful. The production of bovine embryos for commercial purposes follows the steps: they are produced *in vitro* and transferred to synchronized receptors when they reach the blastocyst stage [5].

To their production, the donor cow undergoes a system called *ovum*-pick up to retrieve its oocytes (commercial purpose) or the ovarian follicles could be aspirated on ovaries from abattoir (research purpose) [5].

On contrary to the IVP system, there is a more natural way to produce bovine embryos that is the multiple ovulation and embryo transfer (MOET). It is a treatment used to increase the number of ovulations higher than the average for the species using a sort of hormones and lasting up to 9-10 days [5]. Six to eight days after the artificial insemination of the superstimulated cows in MOET, the embryo presumptively has reach the uterus. In this stage, the embryos are still surrounded by the *zona pellucida* and are tolerant to some handling out of the cow body. Although both *in vitro* (IVP) and *in vivo* (MOET) produced embryos are tolerant, its degree is dependent from the original environment (*in vivo* or *in vitro*) whether them were derived and mainly by their quality [5].

There is some difference between embryos derived from IVP or MOET. The embryos produced *in vivo* have round shape, brownish colour, plasmatic membranes of the blastocysts closely to the *zona pellucida*, and the inner cell mass (ICM) is surrounded by small intercellular spaces [6, 7]. The embryos produced *in vitro* generally are darker (this has been associated with a higher accumulation of lipid or lipid like granules in the cytoplasm due to the use of serum in the culture medium) [8]. Blastomeres of *in vitro* embryos appear more swollen, the perivitelline space is smaller at all pre-compaction stages [8], have a decreased cell count [9] and an increased number of vacuoles in comparison to those *in vivo* produced [10, 11].

### 1.1. Classification of embryos by morphological analysis

The embryonic quality is determinate based on the number and appearance of cells. This classification follows the International Embryo Transfer Society (IETS) [12], using two codes of description, that is, one to assign the quality and the other to the stage of development. The standard evaluation is performed with a stereomicroscope at 50 to 100X magnification. The diameter of bovine embryo is 120 to 190 µm, including the *zona pellucida* (12 to 15 µm). The ideal embryo is compact and spherical with blastomeres of similar size, color and texture uniforms; cytoplasm cannot be granular or vesiculated. The periviteline space should be clear and the *zona pellucida* uniform not cracked or collapsed. The code for the stage of development is numeric, varying from “1” to unfertilized oocyte or 1-cell embryo, to “9” for expanding hatched blastocyst. The code for quality is based on the morphological integrity of embryos, and it is also ranging from “1” (to good or excellent), when they have symmetrical and spherical mass, the cellular material is at 85% intact and irregularities are relatively minor; “2” (to fair) when the embryos have moderate irregularities in the overall shape of embryonic mass or size and color; “3” (to poor) when major irregularities in shape or other parameters are observed; and “4” (to dead or degenerating embryos) that is, a non-viable structure.

Despite this classification, to evaluate bovine embryos is directly affected by the embryologist’s accuracy, experience and mood [13]. The reason for this interference is that the morphological analysis does not measure any objective variables to determine the embryo classification. The reason is that the human vision is subjectively being able to extract numerous information of an image but ignoring some kinds or not observing another one and it is based on a comparison among objects or images. Every time our eyes are like a tool to analysis results of experiments, but when we have to analyze an image, there is a difficulty of at judging color or brightness of shapes and features [14]. Thus the analyses by an embryologist may have low reproducibility [15]. Farin *et al.* [16] described two kinds of error by embryologist that support that affirmation. First is the inter-evaluator error, when the same embryo can be classified with different quality grades by different embryologists or (intra-evaluator error) when the same embryologist classifies the same embryo in different grades. This could specially to occur when the quality grade is borderline, the evaluator is inexperience, he is tired, or his mood is altered. Due all the reasons already mentioned, the embryo evaluation is less reproducible and objective as desired. However, it is one of the most critical steps of the embryo transfer procedure (IVP or MOET) and has a strong relation to a successful pregnancy establishment [12].

### 1.2. The semi-automatized methods to evaluate the embryo quality

Therefore, several methods have been or are being developed to improve the process of embryo classification that has not external effects. Many embryologists select the oocytes/embryos by a non-invasive examination based on simple observation focused on morphology and kinetics of their developmental stage (third day of culture or blastocyst stage) [17]. Among then, embryo metabolism analysis, cellular respiration measurement, evaluation by time-lapse video [18], the quality on *in vitro* growth of embryos, the integrity of blastomeres membrane [18, reviewed by 19; 20] and electron-microscopy analysis [18, reviewed by 21]. Other kinds of evaluation have been used in attempt of find a better solution to classify the embryo quality. As example, Melo *et al*. [22] described an automatic segmentation procedure of bovine embryos, without the use of Artificial Intelligence (AI) and whose proposal was calculated the sensitivity, specificity and accuracy’s method. The database used in this work was made up of 30 histological slides (staining with Sudan Black B) in different stage of development: early cleavage, morula and blastocyst and, the image was acquired by inverted light microscope Olympus IX71 with digital camera coupled (Lumenera Innity 1-1). Each slide was photographed at a magnification of 100x, using 10x objective lens and ocular lens. Although the positive results and the high rates of sensitivity and specificity it is not applicable to the bovine embryos yet.

Time-lapse is a non-invasive technology mainly used to human embryos to measure morphokinetic parameters such as the timing of karyogamy, time intervals between cytokinesis and its abnormal events that results in uneven blastomeres size [23, 24]. With this technology, the embryos can monitored without removing them from the incubator as in another morphological and dynamics analysis. Images are made by a camera inbuilt the incubator that capture images of the embryos at timed intervals [reviewed by 18; 25]. Among the current time-lapse systems, two are the most widely used technologies: the Embryoscope/Fertilitech (an incubator with an integrate time-lapse system) [26, 27] and the Primo Vision/Vitrolife (a compact digital inverted microscope system) [26, 27] and, both use bright field technology.

Other kind of analysis of embryo quality is the semi-automatized grading method of human blastocyst using a support vector machine. It performs the classification by determining a separation rule between two sets of feature values (support vectors) [28]. Among 17 classification methods tested by Meyer [29], this produced the best performance.

From photographs of oocyte and embryos at the 4-cells stage, Manna *et al.* [17] used AI techniques that are based on (*i*) segmentation (selection of the correct region of interest in the image) and pre-processing for reducing artefacts due to noise, blur or illumination conditions; (*ii*) feature extraction (it is usually dependent of numerical descriptors) that represent in a compact way the starting image (such as LBP) [17, 18, 30]; and (*iii*) definition of a classification system, in which the classifier is trained using the data stored in the knowledge base.

Several authors proposed the use of mathematical and statistic tools for the evaluation of the embryo quality, like the multivariate logistic regression with eight predictive factors for the classification of embryos according to implantation potential [18, 31], computer-assisted scoring system (CASS) [18, 32] that is supposed to have a higher discriminatory power for embryo selection, over the standard scoring system that has intrinsic examiner variability. It was also used the multivariate logistic regression (LR) system together with multivariate adaptive regression splines (MARS) that had shown improvement on predictive model when using the computer assisted scoring system associated with data mining. Nevertheless, any of these methods were totally effective in the prediction and the morphological analysis still is widely used to evaluate the embryo quality [18, reviewed by 33].

### 1.3. The Artificial Intelligence technique

The Artificial Intelligence (AI) is the technique that has potential to develop objective, reproducible and non-invasive methodology to predict embryo quality with high levels of accuracy [18]. A specific method as artificial neural network (ANN) along with genetic algorithm (GA) could be used to simulate an accuracy predictive model [34]. ANNs are inspired by the early models of sensory processing by the brain that has a highest and interconnecting neurons network and communicate by electric surge (through axons, synapse and dendrites). An ANN can be creating using a model of neural network in a computer. Applying algorithms that imitate neurons real process we can make the network learn to solve problems. This is make of interactive way presenting examples with known classification, one before other (it is called learning or training) [35]. GA is a search and optimization method inspired by genetic mechanisms under natural evolution [18, 36], whilst the ANN is a technique based on how human neurons transmit and process information to learn and it is indicated for the resolution of complex and nonlinear problems. Once properly trained, the ANN is capable to perform prediction from new data which it has never had access [37, 38]. It was first proposed for mouse embryo (*in vivo* produced) and posteriorly to *in vitro* produced bovine blastocyst but comprising the stages between the initial and expansion [18; reviewed by 39, 40].

Supported by the GA, three ANNs (with the best accuracy) were developed after to have been trained to classify bovine blastocyst images based on the IETS standards [41, 42]. In that study, 482 blastocyst images dataset were used to train some ANNs, from which the best obtained 76.4% of accuracy. Three experienced embryologists from the major Brazilian and world leader company (*In Vitro Brazil*, Mogi Mirim, São Paulo state, Brazil) were responsible to previously evaluate the embryo quality (according IETS standards) to support the ANN learning.

### 1.4. The Blasto3Q algorithm

The work mentioned above, resulted in two different interfaces, one using the MATLAB^®^ platform and an interface using multiplatform approach for online purpose. Both of them, we called “Blasto3Q” program (3Q is refer to the three qualities of blastocysts evaluation) [41, 42]. In the first case, the interface created (in accordance with the recommendations for the commercial use of the program) allows the user interact with the program in a fast and intuitive way. This interface contains a description of the 24 variables that are extracted from the image and analyzed by the ANNs that classifies the same loaded image through the best three obtained ANNs. The second developed interface allows the on-line use of Blasto3Q via a cell phone (smartphone) or PC.

#### 1.4.1. The process to image acquire

Using MATLAB platform, it was possible the automatized analysis without the embryologist’s intervention. For standardization, the software consecutively followed the steps of image import (BMP or JPG), conversion to greyscale (8-bit grayscale), resolution and proportion adjustment (we choose 640×480 pixels as the default resolution because it is the lowest standard and provides sufficient information for interpretation), and intensity adjustment (1% of all information becomes saturated between light and dark pixels; Rocha *et al.*, 2017a, b). These steps facilitate the next process, the segmentation.

#### 1.4.2. Image segmentation

To the image segmentation they were properly isolated from the background and were then subjected to the information extraction techniques. The segmentation algorithm is composed for 4 steps describe below (more details in our previous works) [41, 42]. The first step was calculating the magnitude of the image gradient and the edges were highlighted in all directions, it is important to next steps and essential to characterize the circular shape of the embryo. Following the magnitude gradient, we calculated the binary image and select a value of 128 as intensity threshold. After that, we use the Circular Hough Transform (CHT) to detect the embryo circumference and mapping the image and thus provide an isolated embryo background image [43]. The algorithm search for circles in two stages: first, it searches by circles with radius between 100 and 150 pixels and then, for circles with radius between 150 and 200 pixels to greater accuracy. Therefore, both initial blastocysts (smaller) or expanded blastocysts (larger) can be detected. At the end of both searches, in each image, the detected circle's metrics are compared and the largest radius is used after the best circle is detected. The last step is the blastocyst isolation, where is generated tree versions of the image: (i) the radius of circumference is increase 5 pixels, to make sure that *zona pellucida* is included (called ER); (ii) the radius is decrease by 40 pixel to exclude the trophectoderm selecting the inner mass cells (IMC) and blastocoel for analysis only (called RR); and (iii) the difference between ER and RR. Therefore, only the trophectoderm region was isolated in the image (called TE). The pixel values, which determined the expansion (ER) and the contraction (RR) of the blastocysts images, were obtained by assessment of the image database.

The image texture was defined by repeated random regular patterns in a region of the image that provided information on the surface structure [41, 44]. It is an important characteristic that is used to identify the regions of interest in an image [45]. The statistical methods used to analysis the textures in images was the grey level co-occurrence matrix (GLCM). Considered the most efficient and describes the spatial distribution of the intensity values of the pixels by considering a determined distance and angle, which makes it possible to recognize and classify textures [44, 46].

#### 1.4.3. Extraction of variables

After image standardization and segmentation, 36 variables were extracted, sufficient to extract all the relevant information for the representation of the analyzed bovine blastocyst image. For the complete description of variables, see previous work [42]. The collinearity analysis was performed to eliminate those variables that were correlated with one another (*i.e.*, redundant). It was also possible to determine the Variance Inflation Factor (VIF; which represents the degree of independent variable multi-collinearity when compared with the other independent variables). Collinear variables can be those with higher VIF values than 10. Thirteen iterations were performed until all the variables had a VIF value of less than or equal to 10. Thus, 24 variables remained for use in the ANN.

For the ANN learning process, the Backpropagation algorithm was used [33]. The total of images (n=482) was divided into training (70%), validation (15%) and test (15%). The accuracy of the obtained ANN was verified according to the error between the real values (the mode of the embryologists’ evaluation) and the values obtained by the ANN [47]. Currently, there is no standard method for obtaining the best architecture (the number of neurons in each layer, number of layers, training and transfer functions) of the ANN for a fittest solution of a problem [41, 48]. Thus, the GA technique was used to improve the efficiency on determining the best ANN architecture.

We considered that the population to be studied by the GA was formed by different ANNs architectures, and the goal was to obtain networks that had the lowest error in the blastocyst image classification.

The GA technique developed in our study considered the creation of an initial ANN population with different architectures (each one is called “individuals”), which was randomly generated and composed of 100, 200 or 300 individuals. Each one was defined by a “genetic code”, that is, a specific pattern of 9 different “genes” representing the variables (*i.e.*, the number of neurons in the first, second and third hidden layer; the transfer function for the first, second and third hidden layer; the transfer function for the output layer; the training function to be used; and the number of hidden layers to be used). Each ANN was trained and tested, and their success percentage for the degree of classification was assessed.

This chapter is a continuation of a previous work [41, 49], extending it with new data and aiming the development of a Graphical User Interface to the users that could not purchase or access an inverted microscope to capture digital images. In addition, embryologists from around the word can access the technique online (by the internet), without downloading or installing the software executable. Furthermore, we describe the necessity of smartphone adapters use for stereomicroscope ocular lens to grade blastocysts in a real-time.

## 2. Methodology

Embryos were produced *in vitro* from *cumulus*-oocytes complex from antral follicles of slaughtered cows based on the protocol previously published [50]. On the day 7 of *in vitro* culture (day 0 was on *in vitro* insemination), the embryos were recovered from the culture. Only the embryos morphologically classified as expanded blastocyst with inner cell mass (ICM) relatively smaller than blastocoele and a thin layer of trophectoderm [12], was utilized to evaluation. First, the expanded blastocysts were separated on phosphate buffered saline (PBS) solution to the caption of image. In the experimental group, the embryo digital image was captured with a Motorola smartphone G4 Plus (ANDROID™ 7.0 system) with 16 megapixels of camera, lens aperture of F/ 2,0 and digital zoom of 4x through the ocular lens from a Leica stereomicroscope (S8 APO - 10 x magnification) with maximum zoom magnification (8 x). Different of previous study, it was not necessary the use of an extra macro lens coupled to the smartphone by a clip to capture the images from the stereomicroscope (Figure 1).

**Fig. 1.**
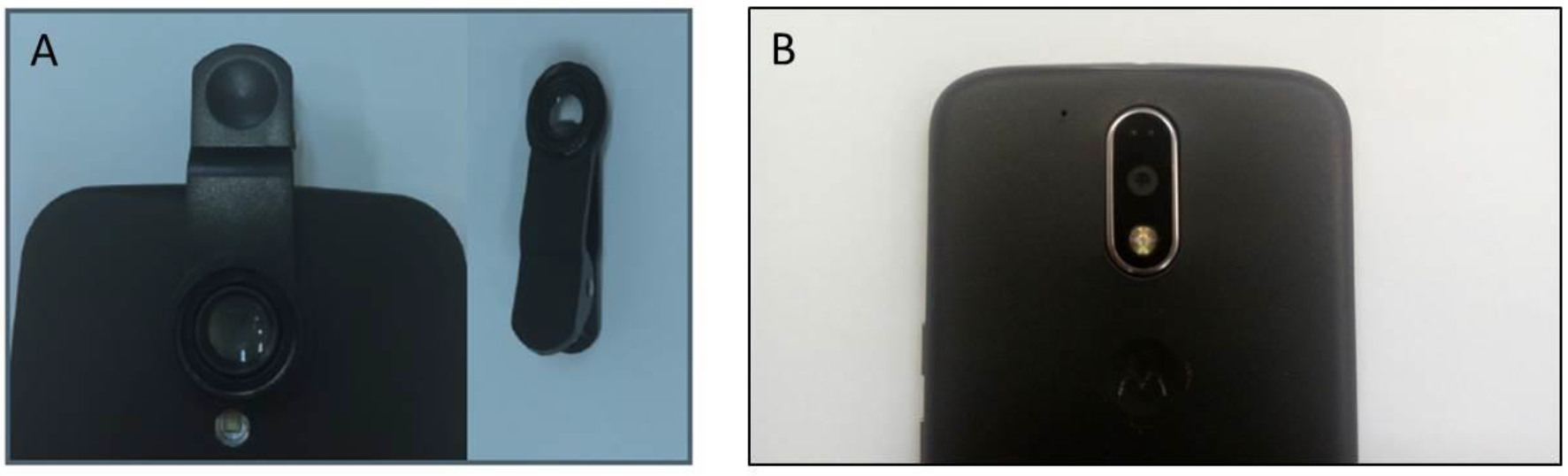
A) Macro lens coupled to the smartphone lens by a clip. Its use was proposed in our previous study (Ciniciato *et al.*, 2017). B) In the present work that device was not necessary since there was no difference (in quality or size of the images acquired by the phone) when the smartphone lens was directly juxtaposed to the stereomicroscope ocular lens.

The images were stored in JPG format. Then, images of the same embryos were also acquired by inverted microscope Nikon Eclipse T*i* coupled to a Nikon Digital Sight DSRi1 at 40 x magnification in the software NIS Elements Ar 3.0 (control group). The images were stored in JPG format in 8-bit color (RGB) at a resolution of 1280 × 1024 pixel. To the capture of pictures with the smartphone and inverted microscope, the embryos were adjusted in a standardized position with ICM perpendicular to the focal plane (Figure 2). Each picture contained only one blastocyst and it is important to note that when it came from the smartphone, the embryo occupied a smaller area of the entire image compared with images from the inverted microscope.

**Fig. 2.**
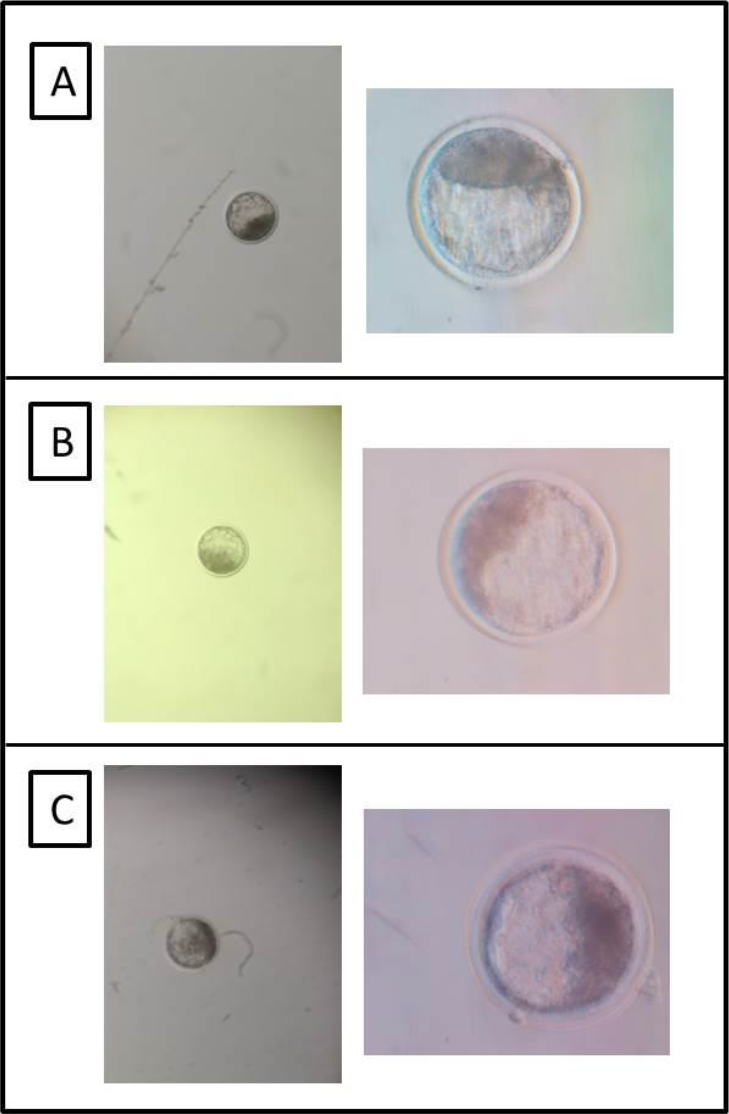
Illustrative samples of the images captured by the smartphone without macro lens (left column) and with the inverted microscope (right column), from three different embryos (lines A, B and C). When the smartphone was used, the embryos occupied a smaller area of the image in comparison to the inverted microscope image.

Embryos from phone images occupied on average 3.01% of total area of the image and their diameter was intercepted, on average, by 21.57% of a line from the extreme points at the largest width of the image (ImageJ; https://imagej.nih.gov/ij/index.html). In control group, the embryos had an average occupation of 26.95% of total area of they were intercepted on average by 49.61% on the maximum width.

To analyze the acquired images, was used the desktop version of Blasto3Q software. First, the image was uploaded on the program, so the image was processed and evaluated by the already trained algorithm Blasto3Q, which is detailed described in [42]. In the software, the uploaded images underwent an automatized segmentation process, whereby the images were properly isolated from the background. The image could be fully, partially or non-segmented (Figure 3). To have a reliable result (*i.e.*, a classification by Blasto3Q), numerical variables should be extracted from the image. Once non-segmented images did not produce those variables, only the images that were properly or partially segmented were considered to the comparison of evaluation performance.

**Fig. 3.**
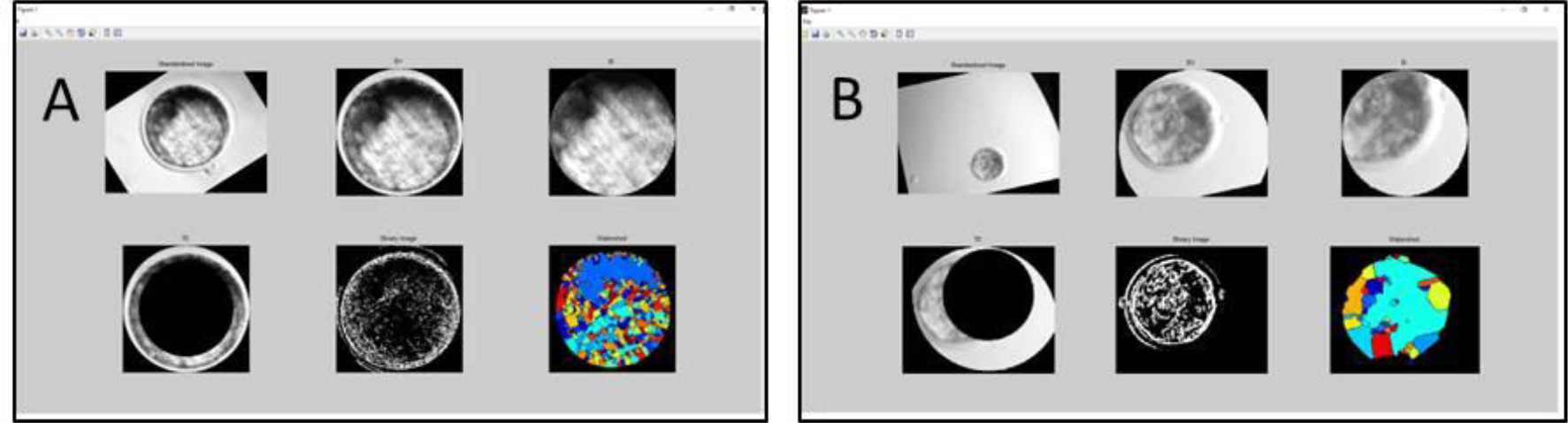
Comparison between perfect (A) and partial segmentation (B) of a blastocyst image. All the images from control group produced a perfect segmentation, whereas in the experimental group there was no perfect segmentation at all images processed.

After the segmentation, the images were submitted to quality grade classification of the three ANNs of the Blasto3Q program to a comparison through the software. For this, one of the ANNs is selected by bottoms at the graphic user interface to analyze the uploaded image. Each ANN deliveries only one quality grade to the user, that is, poor, fair or excellent/good. The output of the same ANN sometimes differed from the same embryo when the sources (control or experimental groups) were different (Figure 4).

**Fig. 4.**
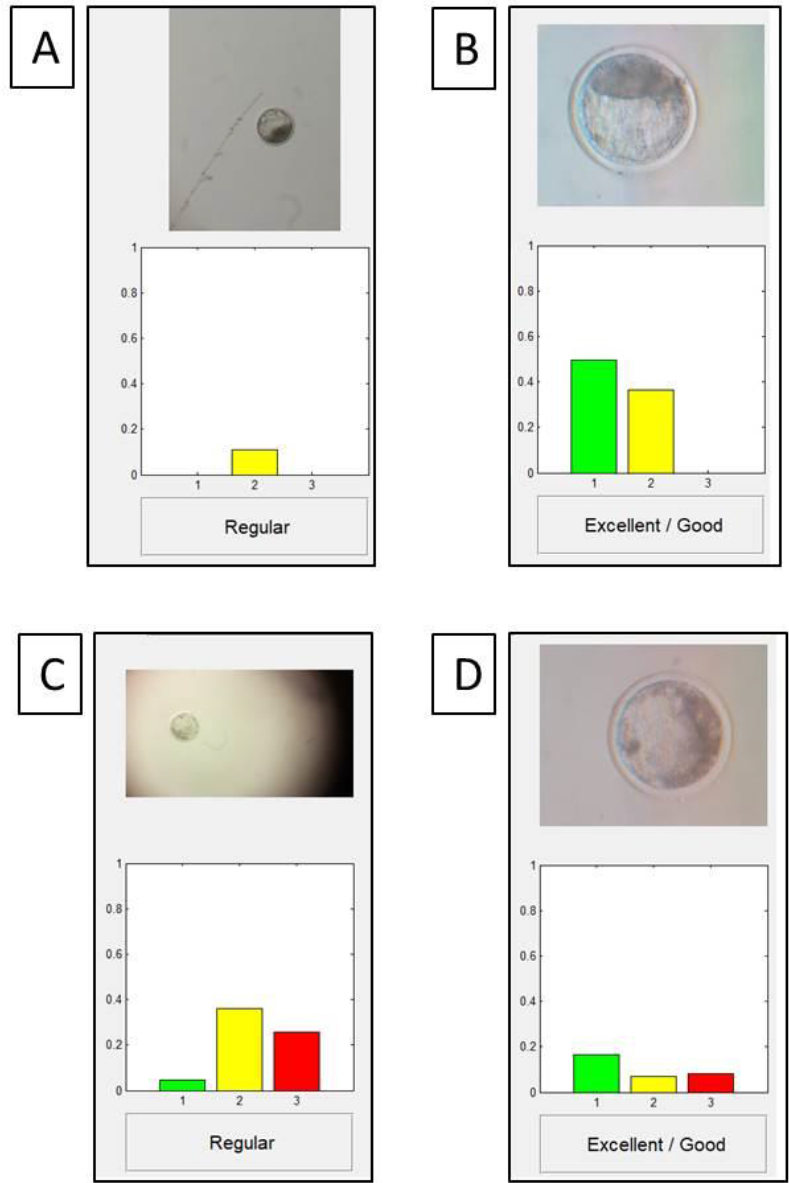
Illustrative samples of the result showed in the graphic user interface (GUI) of Blasto3Q. The GUI shows the uploaded image (raw image) and the classification done by one of the chosen ANNs. Images A and B are from the same embryo when the image was captured by a smartphone (A) and an inverted microscope (B). Images C (smartphone capture) and D (inverted microscope capture) are both from another embryo. The GUI differed between the sources (group) regarding to the showed classification, yet the embryo was the same. The colored bars in the GUI above numbers 1, 2 and 3 - and meaning the quality grades (excellent/good, fair, and poor, respectively) - allow a visual conclusion of what was the higher value (*i.e.*, the highest bar is the quality grade classification).

## Results

Eighteen *in vitro* produced blastocysts were used to obtain distinct images (two biological replicates were performed with 11 and 07 embryos). The same embryo was submitted to digital image capture by the control group (inverted microscope with 40 x of magnification) and to experimental group (stereomicroscope with maximum of magnification [80 x] plus 4 x zoom from the cell phone). The 36 images obtained from control and experimental groups were uploaded on the program Blasto3Q (from an executable file).

Each image from both sources was evaluated for segmentation (*i.e.*, the proper isolation of the embryo and of its parts) since that isolation is a prerequisite to the program obtain numerical values from segmented parts of the image. All images from control group were properly segmented (as showed in the Figure 3A) whereas none image was in the experimental group. In this, 38.9% of the images (07/18) could not be segmented at all, thus no numerical variables were extracted from the images and no quality grade of the embryo could be evaluated by the Blasto3Q. Eleven images from experimental group (61.1%; 11/18) were partially segmented, *i.e.* although the embryo was isolated and numerical values could be extracted there was a mismatch and some parts of the embryo were lost during segmentation steps (as exemplified in the Figure 3B).

The 11 blastocyst images segmented properly (control group) or partially (experimental group) were submitted to the quality grade classification by the three artificial neural networks (ANNs) of the Blasto3Q program. It is mean that each of the image was evaluated by three different networks providing 33 quality grades for control and experimental groups. The percentage of agreement was calculated between the same blastocyst been evaluated by the same artificial neural network from the two sources (control and experimental groups). A total of 36.4% of agreement (12/33) was obtained by ANNs 1, 2 and 3 (3, 4 and 5 of agreement, respectively). The disagreement (63.6%; 21/33) between the groups were 8, 7 and 6 respectively for ANNs 1, 2 and 3.

Overall, from the 54 potential evaluations of the three ANNs from the 18 images of the experimental group, only 22.2% were concordant with the control (12/54). The remaining 42 evaluations were unable to be performed (21) or were wrongly processed (21).

## Discussion

The alternative source of image capture (from a smartphone) produced a kind of embryo characterization that impaired the Blasto3Q program to properly classify it. It was notorious by the fully segmentation criterion. Since the program was trained with the higher percentage and interception of the embryo image covering the total area of the picture (from an inverted microscope capture) and, consequently, with a higher diameter of the embryo in the image to be recognizable by the segmentation process. We assume that the total failure from the 18 images (experimental group) to be properly segmented was due the smaller area and interception of the embryo image into the total picture uploaded.

As ANN is based on the machine learning concept, the Blasto3Q algorithm learned to classify blastocyst from the pattern observed in the inverted microscope capture. In this way, the 76% accuracy obtained in its training was lost because the evaluation is based from the numerical values extracted of the segmented image. As observed, none image could be fully segmented and only 61.1% were partially segmented in the experimental group. The numerical variables extracted from partially segmentation surely produced a mismatch to the real information contained in the full embryo image. In this way, the bad quality variables extracted (experimental group) were unable to pair the quality evaluation from control group, with only 36.4% of agreed classification between them.

We did not test it, but our hypothesis is that a simple readjustment of the *radius* size to search for the embryo (adjusted for a higher *radius* in the program Blasto3Q, since it was based entirely from inverted microscope images) could improve or even repair the fault observed in this work. The *radius* size is a parameter used by Hough transform to find a circularity in the total image and it is the way to segment only the embryo (by its almost perfect and constant circularity).

The other inferred possibility to correct the error, could be the full re-training of the algorithm and, in this time, with a full database composed only by smartphone blastocyst pictures. Maybe the association of a new phone image database and resizing the radius of the embryo could properly fit the algorithm Blasto3Q to the alternative use of smartphones as the tool to directly capture images from stereomicroscope.

## Conclusion

We concluded that, with the current training of the Blasto3Q algorithm, it was not possible the use of a smartphone camera to directly capture blastocyst images from the stereomicroscope ocular lens. The agreement rate of the embryo classification between the standard and the alternative proposal was too low (22.2%) to be considered feasible.

## Acknowledgements

The author’s research is supported by grants #2012/50533-2, 2013-05083-1, 2006/06491-2, 2011/06179-7, 2012/20110-2 and 2016/19004-4 from São Paulo Research Foundation (FAPESP). We also thank Agência UNESP de Inovação (AUIN) for processing the national and international patents of the invention.

